# Comprehensive Analysis of Peptide-Coding Genes Identifies an LRR-only Microprotein That Regulates Reproductive Organs in *Marchantia polymorpha*

**DOI:** 10.1101/2022.09.27.509623

**Authors:** Haruaki Kobayashi, Shigeo S. Sugano, Kentaro Tamura, Yoshito Oka, Tomonao Matsushita, Tomoo Shimada

## Abstract

In the past two decades, many plant peptides have been found to play crucial roles in various biological events by mediating cell-to-cell communications. However, a large number of small open reading frames (sORFs) or short genes capable of encoding peptides remain uncharacterized. In this study, we examined several candidate genes for peptides conserved between two model plants: *Arabidopsis thaliana* and *Marchantia polymorpha*. We examined the expression pattern in *M. polymorpha* and subcellular localization using a transient assay with *Nicotiana benthamiana*. We found that one candidate, *MpSGF10B*, was expressed in meristems, gemma cups, and male reproductive organs called antheridiophores. MpSGF10B has an N-terminal signal peptide followed by two leucine-rich repeat (LRR) domains and was secreted to the extracellular region in *N. benthamiana* and *M. polymorpha*. Compared with the wild type, two independent knockout mutants had an increased number of antheridiophores, which emerged from meristems. It was revealed in gene ontology enrichment analysis that *MpSGF10B* was significantly co-expressed with genes related to cell cycle and development. These results suggest that MpSGF10B may regulate the reproductive development in *M. polymorpha*. Our research should shed light on the unknown role of LRR-only proteins in land plants.

## 2 Introduction

Over the last two decades, many biologically active peptides have been identified in land plant species that play critical roles in plant growth and development through molecular genetic studies combined with biochemical analyses. Plant peptides are generally defined as less than 100 amino acids in length. Their structural characteristics can be classified into two major groups (Matsubayashi, 2014; Hirakawa et al., 2017; Stührwohldt and Schaller, 2018). One group is small post-translationally modified peptides characterized by the presence of post-translational modifications and by their small size after proteolytic processing (approximately 5–20 amino acids). The other group comprises cysteine-rich peptides (approximately 40–120 amino acids), characterized by an even number of cysteine residues forming intramolecular disulfide bonds. These plant peptides are often generated by proteolytic processing from larger precursors with N-terminal secretion signal sequences and post-translational modifications. Mature peptides generally function as extracellular ligands in cell-to-cell or long-distance signaling by interacting with their corresponding receptors, leucine-rich repeat receptor-like kinases/proteins (LRR-RLK/RLP), on the plasma membrane of target cells.

Since the discovery of systemin in *Solanum lycopersicum* (McGurl et al., 1992), several classes of small secreted peptides have been identified in land plant species. Hormone-like peptides are classified into several families, including CLAVATA3/Embryo surrounding region-related (CLE), inflorescence deficient in abscission (IDA), epidermal patterning factor (EPF), and rapid alkalinization factor (RALF) (Gancheva et al., 2019). These peptide families and their receptors are conserved in embryophytes but are rare in green algae (Ghorbani et al., 2015; Bowman et al., 2017; Olsson et al., 2018), indicating that these peptide-signaling pathways evolved in a common ancestor of land plants and served some advantage in terrestrialization (Furumizu et al., 2018; Moody, 2019). In one example, CLE peptides and their receptors are conserved between *Marchantia polymorpha* and *Arabidopsis thaliana* and regulate the size of meristems (Fletcher et al., 1999; Hirakawa et al., 2019, 2020). Many peptide members are encoded as multiple paralogs in most land plant species, which makes it challenging to analyze their characteristics genetically, while *M. polymorpha* has low redundancy in peptide families (Bowman et al., 2017). Furthermore, *M. polymorpha* provides a feasible system for studying small peptides reverse-genetically: for example, gateway binary vectors for *M. polymorpha* (Ishizaki et al., 2015), agrobacterium-mediated transformation (Kubota et al., 2013; Tsuboyama and Kodama, 2018), and CRISPR/Cas9–based genome editing (Sugano et al., 2018).

Many peptides have been identified with a similarity search for orthologs and forward-genetics, such as *de novo* identification with mass spectrometry (Fan et al., 2022). Although the importance of bioactive peptides in plant cells has been recognized, reverse genetic analysis to search for new peptides has been difficult. This is due to the short sequence length of peptides, which is often judged as noise rather than a gene by conventional gene prediction (Hellens et al., 2016). In plants, an attempt was made to comprehensively search for small open reading frames (sORFs) that are likely to encode peptides. This was made by a powerful gene prediction method in which statistical processing based on the base composition of known protein-coding genes and the transcriptome are combined and a high rate of false-positive prediction is overcome (Hanada et al., 2010, 2013). This study identified 7,901 sORFs as candidates for new genes in *A. thaliana*. Among these sORFs, 49 were found to cause morphological changes when overexpressed (Hanada et al., 2013). Several databases for sORFs have recently been established (Hazarika et al., 2017; Chen et al., 2020; Liang et al., 2021). Furthermore, in the Arabidopsis Information Resource (TAIR) database, small coding genes of less than 120 amino acids in length have increased in number with version updates (Takahashi et al., 2019). These findings suggest that not a few functional sORFs or peptides remain uncharacterized in genomes that are involved in various physiological phenomena.

Here, we searched for novel peptide-coding genes conserved between land-plant species and analyzed their characteristics using the high-quality dataset of *A. thaliana* and the feasible platform for reverse genetics of *M. polymorpha*.

## 3 Results

### 3.1 Selection of putative peptide-coding genes conserved between *M. polymorpha* and *A. thaliana*

We searched genes in *M. polymorpha* that had high sequence similarity to the sORFs in *A. thaliana* to find small genes that are evolutionarily conserved among land plants (datasets of the sORFs are available from https://rdr.kuleuven.be/dataset.xhtml?persistentId=doi:10.48804/SP9WIK). The workflow is shown in Figure 1A. From 24,751 proteins in the *M. polymorpha* dataset, we first selected 6,815 small proteins with a length of < 200 amino acids. We then performed a BLASTp search using this small protein dataset as a database and the translated sORFs dataset in *A. thaliana* as a query. After that, we selected 28 genes in *M. polymorpha* encoded putative bioactive peptides (E-value < 0.0001). The search results are listed (Table 1 and Supplementary Table S1). The resulting genes can be classified into 18 groups (hereafter referred to as “small gene families”; SGF1–SGF18). Among these 18 SGFs, we focused on 7 SGFs, whose functions cannot be presumed from annotations registered in MarpolBase (https://marchantia.info/; version 5), as candidates without characterized functions (Figure 1B, Table 1, and Supplementary Table S1). The sequence alignments between the SGFs in *M. polymorpha* (MpSGF) and the sORFs in *A. thaliana* are shown in Figure S1. They had two patterns: 1) enrichment of identical residues in a narrow range in SGF6 and SGF10, and 2) scattered matches in SGF13, SGF17, and SGF18 (Supplementary Figure S1).

**Figure 1.**
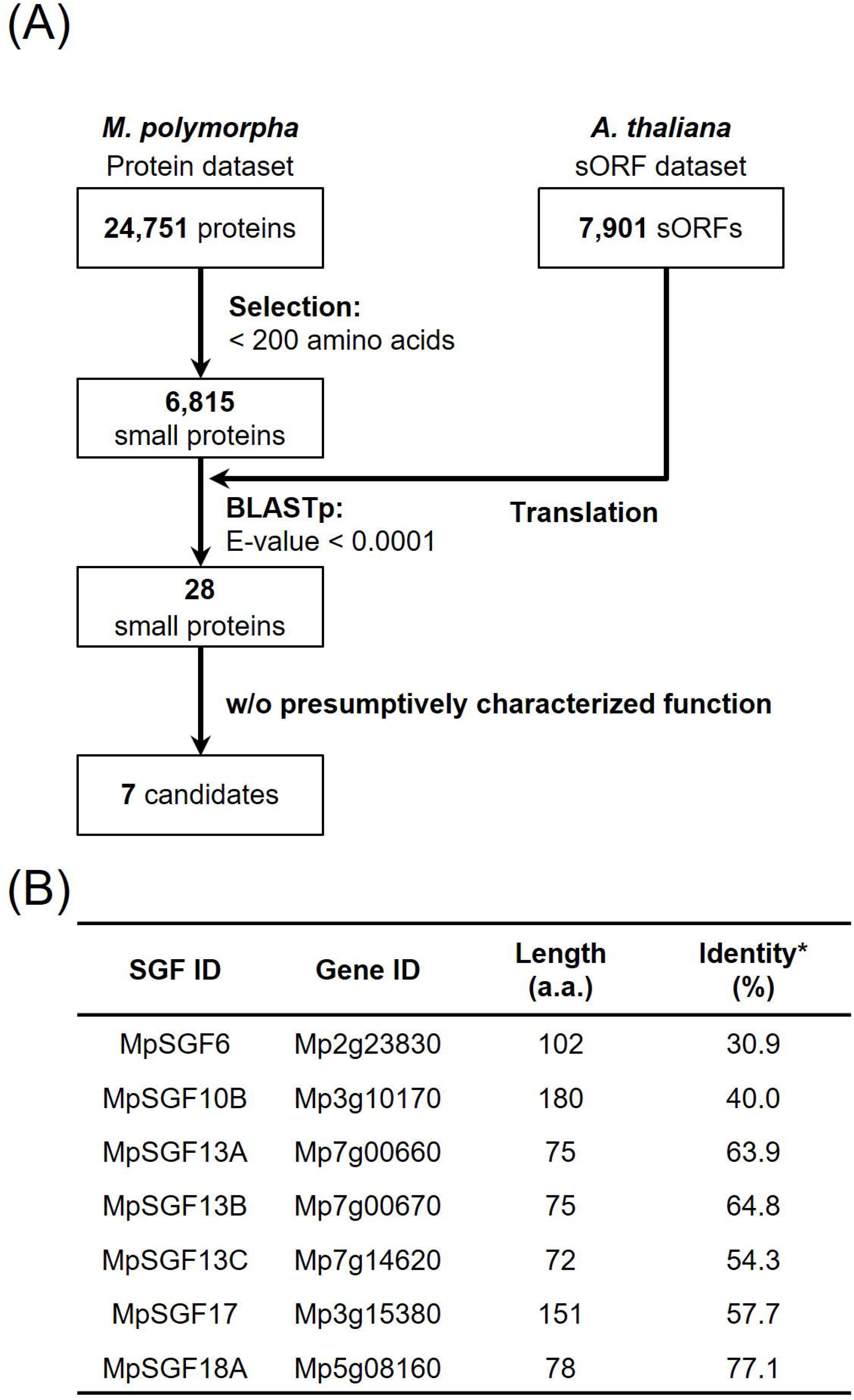
Selection of candidates for *M. polymorpha* peptide-coding genes. **(A)** A pipeline for the selection of peptide candidates. The *A. thaliana* sORF dataset is used as a query for BLASTp analysis. **(B)** A list of selected peptide candidates in *M. polymorpha*. Percentages of identity between MpSGFs and corresponding sORFs in *A. thaliana* were shown.

**Table 1.**
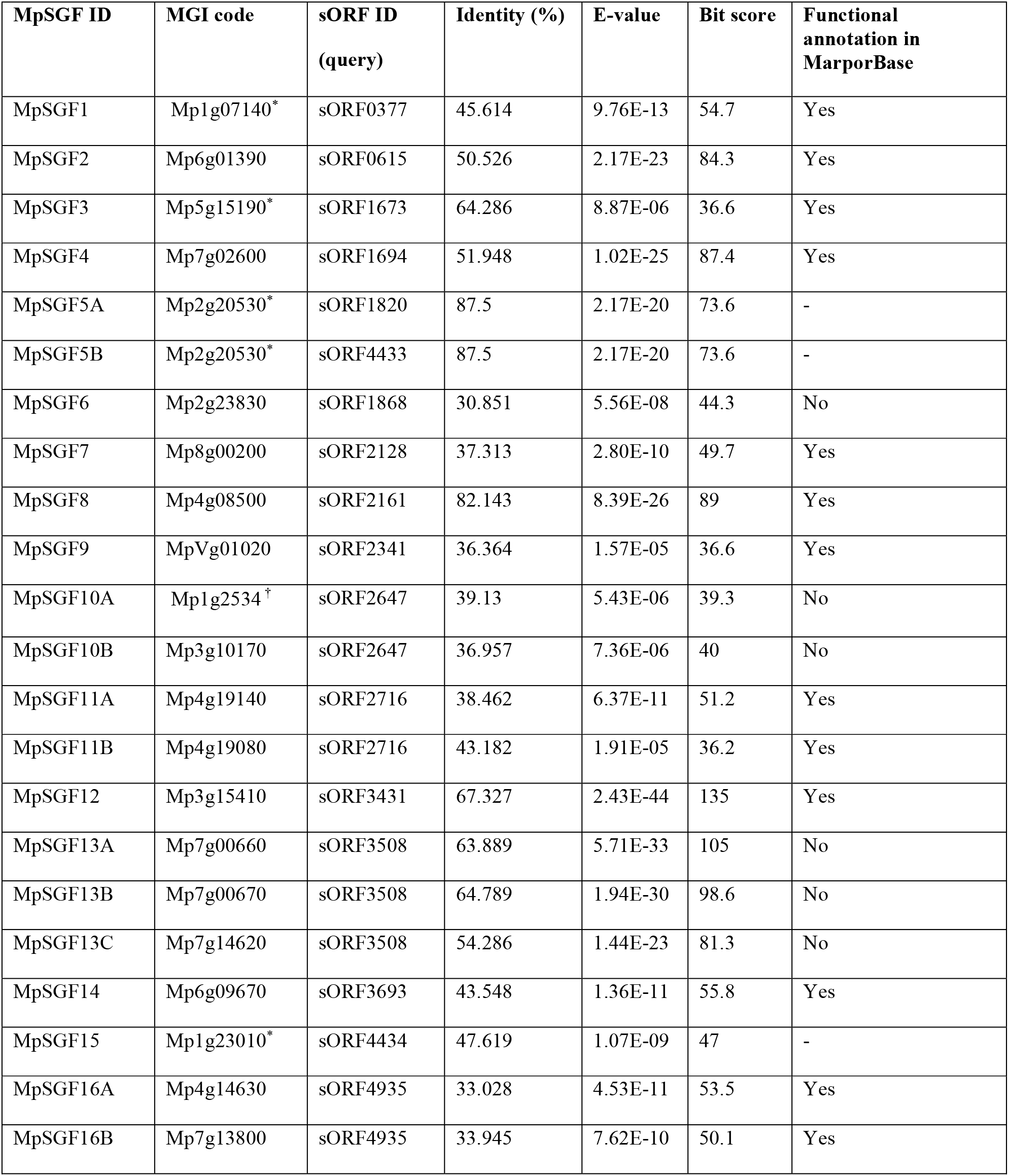
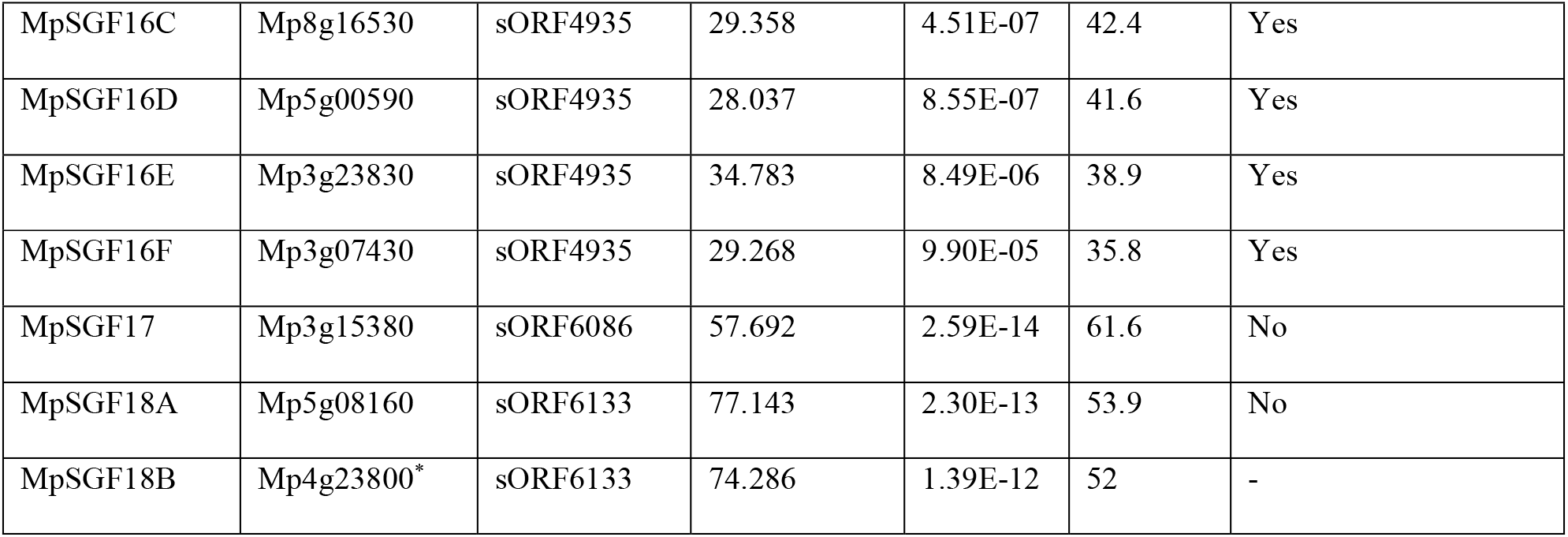
A list of MpSGFs, containing BLAST results and annotations registered in the database (MarpolBase). See Table S1 for detailed information. * Deleted in newer database ver. 6 (released on 2021.3.15). ^†^ non-canonical protein-coding gene (e.g., non-AUG start codon).

| MpSGF ID | MGI code | sORF ID<br>(query) | Identity (%) | E-value | Bit score | Functional<br>annotation in<br>MarpolBase |
| --- | --- | --- | --- | --- | --- | --- |
| MpSGF1 | Mp1g07140* | sORF0377 | 45.614 | 9.76E-13 | 54.7 | Yes |
| MpSGF2 | Mp6g01390 | sORF0615 | 50.526 | 2.17E-23 | 84.3 | Yes |
| MpSGF3 | Mp5g15190* | sORF1673 | 64.286 | 8.87E-06 | 36.6 | Yes |
| MpSGF4 | Mp7g02600 | sORF1694 | 51.948 | 1.02E-25 | 87.4 | Yes |
| MpSGF5A | Mp2g20530* | sORF1820 | 87.5 | 2.17E-20 | 73.6 | - |
| MpSGF5B | Mp2g20530* | sORF4433 | 87.5 | 2.17E-20 | 73.6 | - |
| MpSGF6 | Mp2g23830 | sORF1868 | 30.851 | 5.56E-08 | 44.3 | No |
| MpSGF7 | Mp8g00200 | sORF2128 | 37.313 | 2.80E-10 | 49.7 | Yes |
| MpSGF8 | Mp4g08500 | sORF2161 | 82.143 | 8.39E-26 | 89 | Yes |
| MpSGF9 | MpVg01020 | sORF2341 | 36.364 | 1.57E-05 | 36.6 | Yes |
| MpSGF10A | Mp1g2534† | sORF2647 | 39.13 | 5.43E-06 | 39.3 | No |
| MpSGF10B | Mp3g10170 | sORF2647 | 36.957 | 7.36E-06 | 40 | No |
| MpSGF11A | Mp4g19140 | sORF2716 | 38.462 | 6.37E-11 | 51.2 | Yes |
| MpSGF11B | Mp4g19080 | sORF2716 | 43.182 | 1.91E-05 | 36.2 | Yes |
| MpSGF12 | Mp3g15410 | sORF3431 | 67.327 | 2.43E-44 | 135 | Yes |
| MpSGF13A | Mp7g00660 | sORF3508 | 63.889 | 5.71E-33 | 105 | No |
| MpSGF13B | Mp7g00670 | sORF3508 | 64.789 | 1.94E-30 | 98.6 | No |
| MpSGF13C | Mp7g14620 | sORF3508 | 54.286 | 1.44E-23 | 81.3 | No |
| MpSGF14 | Mp6g09670 | sORF3693 | 43.548 | 1.36E-11 | 55.8 | Yes |
| MpSGF15 | Mp1g23010* | sORF4434 | 47.619 | 1.07E-09 | 47 | - |
| MpSGF16A | Mp4g14630 | sORF4935 | 33.028 | 4.53E-11 | 53.5 | Yes |
| MpSGF16B | Mp7g13800 | sORF4935 | 33.945 | 7.62E-10 | 50.1 | Yes |
| MpSGF16C | Mp8g16530 | sORF4935 | 29.358 | 4.51E-07 | 42.4 | Yes |
| MpSGF16D | Mp5g00590 | sORF4935 | 28.037 | 8.55E-07 | 41.6 | Yes |
| MpSGF16E | Mp3g23830 | sORF4935 | 34.783 | 8.49E-06 | 38.9 | Yes |
| MpSGF16F | Mp3g07430 | sORF4935 | 29.268 | 9.90E-05 | 35.8 | Yes |
| MpSGF17 | Mp3g15380 | sORF6086 | 57.692 | 2.59E-14 | 61.6 | No |
| MpSGF18A | Mp5g08160 | sORF6133 | 77.143 | 2.30E-13 | 53.9 | No |
| MpSGF18B | Mp4g23800* | sORF6133 | 74.286 | 1.39E-12 | 52 | - |

### 3.2 Expression analysis and subcellular localization of MpSGFs

Next, we examined the expression pattern of the selected 7 MpSGF genes. It was revealed in reverse transcription-PCR (RT-PCR) analyses that the expression of all MpSGF genes was detected in thalli and antheridiophores, corresponding to vegetative organs and reproductive organs, respectively. These results indicate that these MpSGFs function in both vegetative and reproductive states in *M. polymorpha* (Figure 2A). The results of RT-PCR were consistent with public transcriptome data (Bowman et al., 2017). However, the expression levels of each gene varied between organs (Supplementary Figure S2).

**Figure 2.**
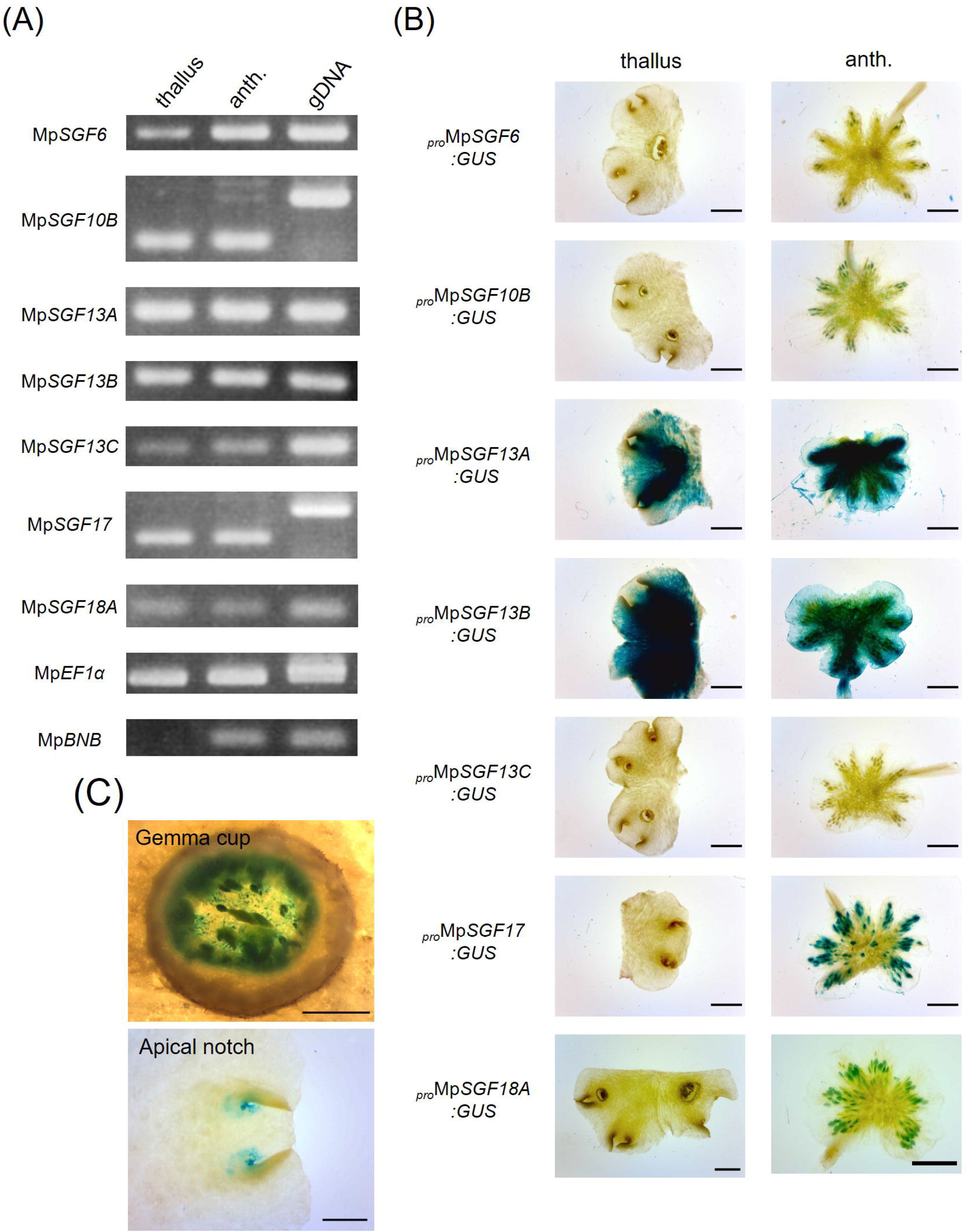
Expression patterns of MpSGFs in *M. polymorpha*. **(A)** RT-PCR in 2-week-old thallus (thallus) and antheridiophore (anth.). *MpEF1α* is used as an internal control. *MpBNB* is used as a marker gene for antheridiophore (Yamaoka et al., 2018). **(B)** Representative images of the GUS reporter assay for expression patterns of MpSGFs. Two-week-old thalli and antheridiophores were observed for the visualization of promoter activities. Bars = 2 mm. **(C)** Magnified images of GUS signal in the bottom of gemma cups and apical notches of *_pro_MpSGF10B:GUS* plants. Bars = 0.5 mm.

To investigate the spatial distribution of the selected MpSGFs in planta, we observed their promoter activities using transgenic plants harboring the ß-glucuronidase (GUS) reporter. We introduced the GUS reporter constructs with 4–5 kb upstream regions of coding sequence (CDS) into the male wild type plant Takaragaike-1 (MpTak-1). Two-week-old thalli and antheridiophores were sampled and stained as vegetative and reproductive organs, respectively (Figure 2B). The GUS activities of the promoter of MpSGF6, MpSGF17, and MpSGF18 were observed in antheridiophores, especially around antheridia, but not in thalli. Intense GUS activities were detected in antheridiophores and thalli in the MpSGF13A and MpSGF13B lines, while no GUS activity in the MpSGF13C lines. In the MpSGF10B line, GUS activity was specifically observed around apical notches and bottoms of gemma cups, in addition to in antheridiophores (Figure 2C).

We performed a transient expression assay for the yellow fluorescent protein MpSGF (MpSGF-YFP) fusion proteins using *Nicotiana benthamiana* to determine the subcellular distribution of MpSGFs. The YFP signal of MpSGF6-YFP was co-localized with chlorophyll (Figure 3A and Supplementary Figures S4A,C). While MpSGF10B-YFP was detected in the periphery of epidermal cells and appeared to overlap with the plasma membrane, the signal was visualized in an extracellular region after plasmolysis treatment (Figures 3B,C), suggesting that MpSGF10B can be transported to the extracellular region. The YFP fluorescence of MpSGF13A, MpSGF13B, and MpSGF13C was ambiguously observed in the cytosolic compartment (Figures 3D,E,F). The YFP signal of MpSGF17-YFP was co-localized with chlorophyll (Figure 3G and Supplementary Figure S4,D). MpSGF18A-YFP was observed in a cytosolic compartment (Figure 3H).

**Figure 3.**
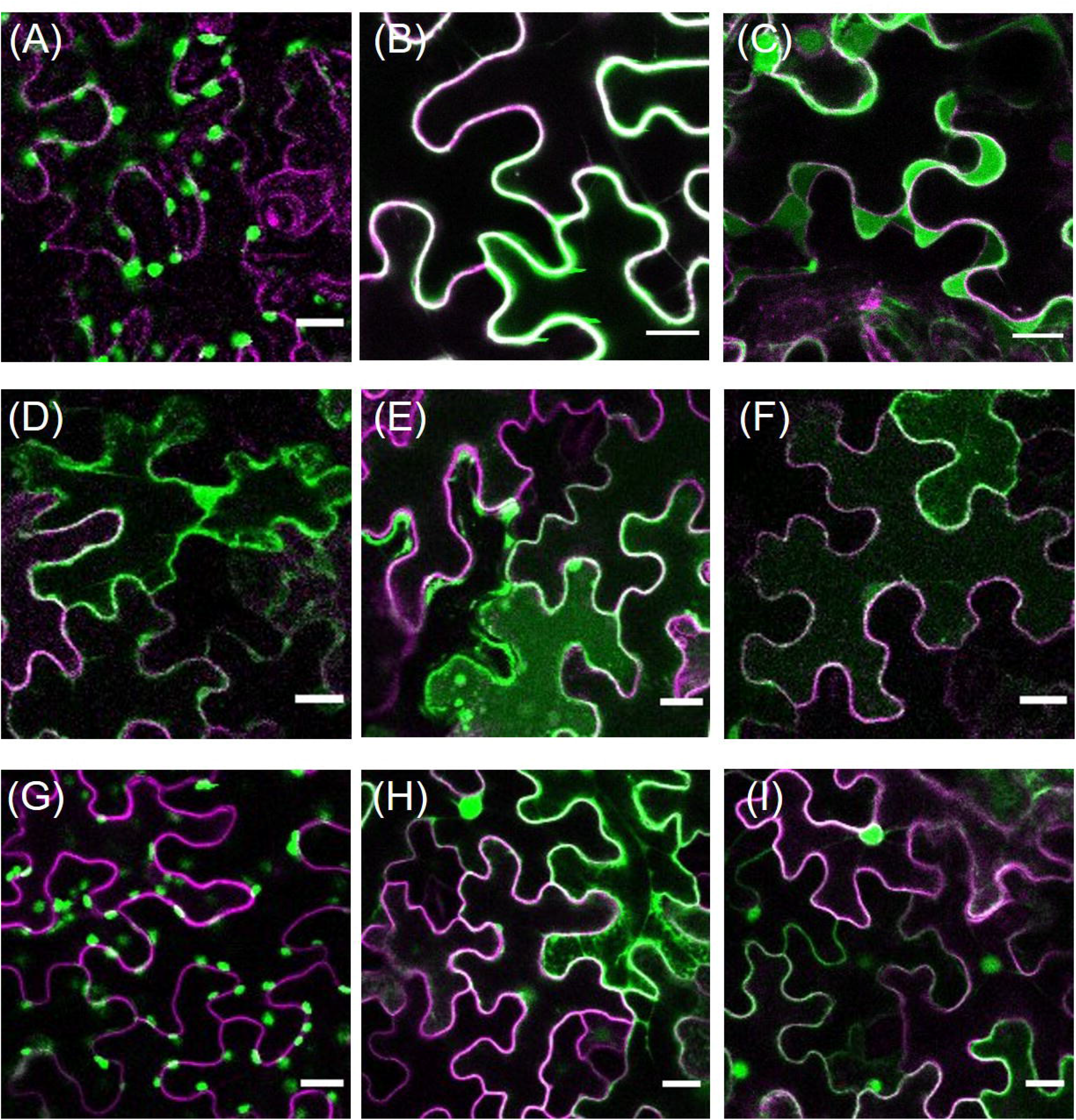
Subcellular localization of MpSGFs in *N. benthamiana*. Confocal fluorescence microscopy images of MpSGF-YFP transiently expressed in *N. benthamiana;* **(A)** MpSGF6, **(B)** and **(C)** MpSGF10B under isotonic and hypertonic conditions, respectively, **(D)** MpSGF13A, **(E)** MpSGF13B, **(F)** MpSGF13C, **(G)** MpSGF17, **(H)** MpSGF18A, and **(I)** YFP only. The plasma membrane, which was visualized by RFP-LTI6b, and YFP were shown in magenta and green, respectively. Bars = 20 μm.

### 3.3 *MpSGF10B* may affect reproductive induction and cell division activity

We confirmed that the C-terminal fusion of YFP with MpSGF10B was also secreted to the extracellular region in *M. polymorpha* (Figure 4A). Thus, we focused on MpSGF10B as a candidate for a secreted bioactive peptide, and further analyses were performed. *MpSGF10B* encodes a protein with an N-terminal signal peptide, followed by two leucine-rich repeats (Figure 4B). We generated *Mpsgf10b^ge^* mutants using the CRISPR/Cas9 system to examine the function of the MpSGF10B in *M. polymorpha. Mpsgf10b-1^ge^* and Mpsgf1*0b-2^ge^* harbor frameshift mutations that generate early stop codons after the predicted signal peptide sequence and in the middle of the sequence, respectively (Figure 4B). In both *Mpsgf10b^ge^* mutants, the number of antheridiophore was increased compared to the wild type (Figure 4C). These results indicate that *MpSGF10B* may repress reproductive induction in liverwort.

**Figure 4.**
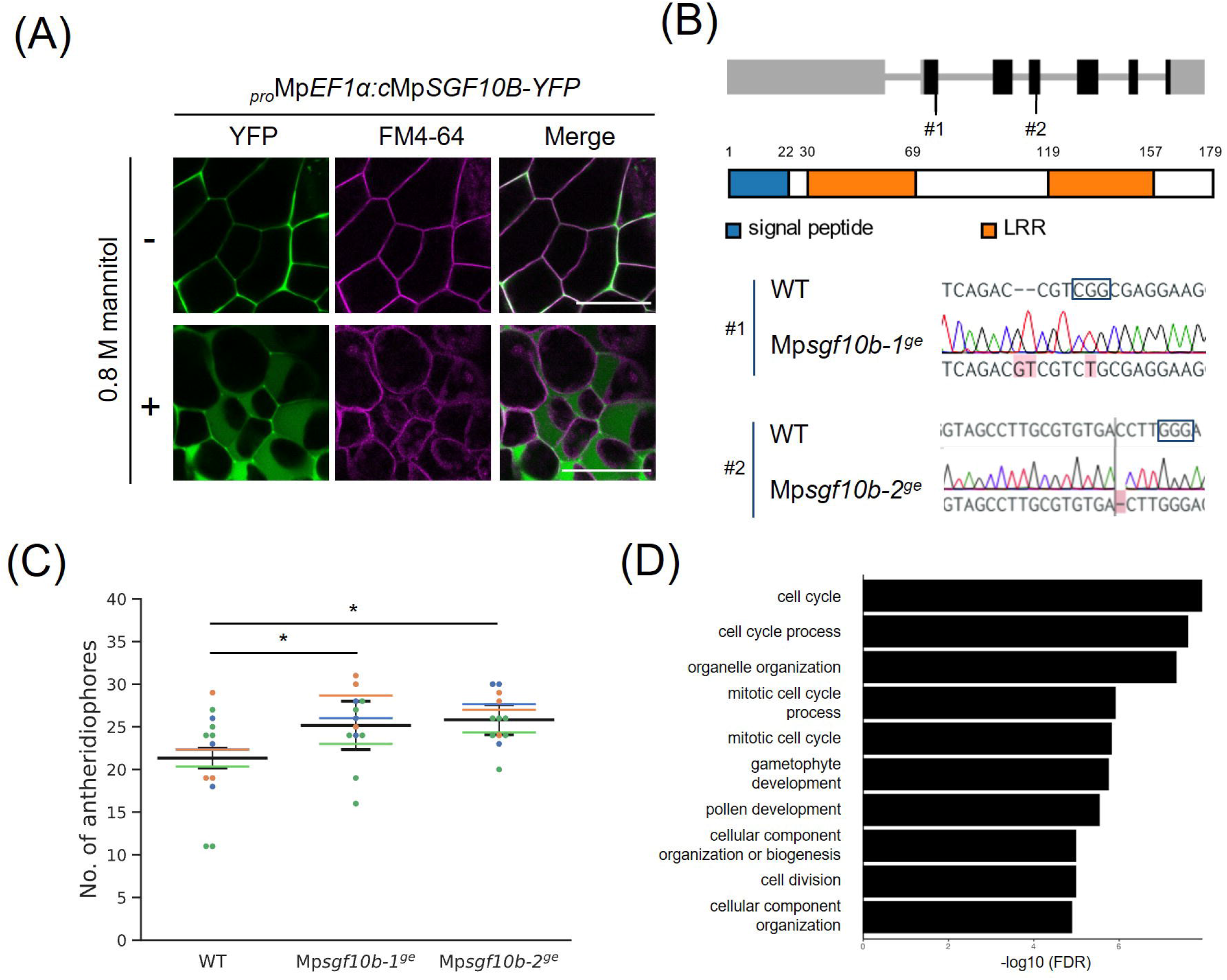
Reverse genetic analysis of MpSGF10B. **(A)** Subcellular localization of MpSGF10B-YFP constitutively expressed in *M. polymorpha*. Three-day-old gemmalings were observed. The plasma membrane was visualized with FM4-64 (magenta). Shown data are representative ones from 5 independent transgenic lines. Bars = 50 μm. **(B)** Gene structures and genotypes of *MpSGF10B* knockout mutants using CRISPR/Cas9. (top) Structure of MpSGF10B/Mp3g10170 locus with the positions of the designed guide RNAs (#1 and #2). Black boxes, gray lines, and gray boxes show coding exons, introns, and untranslated regions. (Middle) The predicted product of MpSGF10B is shown. (Lower) Sequences of genome editing alleles are shown. The PAM sequence is shown in enclosed characters. Edited bases are indicated with red shades. **(C)** The number of antheridiophores in an individual plant. *n* = 3, 3, and 5 for all lines from three independent experiments, respectively (indicated in distinct colors). Horizontal lines indicate the average numbers grouped by the experiments. Black lines show the total means ± SD. \**P* < 0.05, Dunnette’s *t*-test against wild type (WT). **(D)** GO enrichment analysis of highly co-expressed genes with MpSGF10B. The 10 high-ranking biological processes are shown at a 1% false discovery rate (FDR).

Next, we examined the functional categories of co-expressed genes with *MpSGF10B* by gene ontology (GO) to infer the functional categories of *MpSGF10B* at the molecular level. We calculated the R-values (Pearson’s coefficient of correlation) of expression intensities between *MpSGF10B* and all annotated genes in various organs and conditions. We then defined those annotated genes with the 1% high-ranking R-values as co-expressed genes (Hanada et al., 2013). The over-represented functions of the co-expressed genes were related to cell cycle and development at lower FDR values among the GO categories of biological processes (Figure 4D and Supplementary Table S2). This was consistent with the abovementioned reporter assay, since the promoter activity of *MpSGF10B* was detected around the apice and the bottom of gemma cups, in which cell division was taking place for the growth and production of gemmae, respectively (Figure 2C). Collectively, these data suggest that MpSGF10B may function as a regulatory factor of the cell cycle or cell division at the molecular level.

## 4 Discussion

Over the past few decades, various peptide-coding genes have been predicted in *A. thaliana* (Hanada et al., 2007, 2013). Since many bioactive peptides tend to be conserved among land plant species, it is reasonable that bioactive peptides can be detected by comparing evolutionally distant plant species (e.g., bryophytes and flowering plants). We investigated novel peptide-coding genes conserved between *A. thaliana* and *M. polymorpha* through a BLAST-based similarity search followed by expression and loss-of-function analyses in this study. Finally, we identified a gene, MpSGF10B, encoding a secreted LRR-only protein that may be involved in developing reproductive organs in *M. polymorpha*.

### 4.1 MpSGF10B encodes an extracellular LRR-only protein

We showed that the LRR-containing protein MpSGF10B was secreted to the extracellular region in this study (Figures 3B,C). LRR proteins in plants act in various biological processes. According to the subcellular localization and structure, LRR proteins are classified into three major groups: LRR-RLKs/RLPs at the plasma membrane, intracellular nucleotide-binding site (NBS)- LRRs, and extracellular LRR-only proteins (Wang et al., 2011). MpSGF10B may belong to the extracellular LRR-only protein group.

LRR-RLKs/RLPs and NBS-LRRs have been well studied and act in plant immune responses, such as various receptors and R-genes (Dievart et al., 2020; Sett et al., 2022). In contrast, extracellular LRR-only proteins have been poorly reported. A well-known LRR-only protein, polygalacturonase inhibitor 1 (PGIP1), participates in disease resistance by inhibiting the activity of fungal pectin-degrading enzymes (Cheng et al., 2021). *Arabidopsis* PGIPs have an N-terminal signal peptide and LRR motifs (PGIP1: Q9M5J9, PGIP2: Q9M5J8 in the UniProt database). Therefore, PGIPs and MpSGF10B have similar features. However, there are some differences between MpSGF10B and rice PGIPs, which are anchored to the plasma membrane or cell wall (Wang et al., 2015) and possess as many as 10 repeats of the LRR motif. It is reported that canonical LRR-only proteins contained 7—13 repeats (Wang et al., 2011). Since MpSGF10B has fewer LRR motifs and is smaller than canonical LRR proteins, it can be considered a non-canonical LRR-only protein (Figure 4B). As such non-canonical LRR-only proteins, NTCD4 (F4JNB0 in the UniProt) and LRRop-1 (Q56X33), have been reported in *A. thaliana*. NTCD4 consists of 153 amino acids and interacts with MoNLP1, a type of microbe-associated molecular pattern, to induce cell death. NTCD4 is secreted in the extracellular region (Chen et al., 2021). LRRop-1, an LRR-only protein of 218 amino acids, has been suggested to contribute to ABA signaling and is localized in the ER (Ravindran et al., 2020).

Several repeats of LRR motifs, which are generally involved in protein-protein interaction, form a twisted super-helical or a small arc-shaped structure (Kobe and Kajava, 2001; Chakraborty et al., 2019). Therefore, it is unlikely that a small number of LRR units can function as canonical receptors. Further studies are necessary to verify the molecular function of MpSGF10B, whether it acts as a coreceptor that assists other receptors or a ligand for signal transduction or serves an entirely different function.

### 4.2 MpSGF10B might regulate the development of reproductive organs through auxin signaling in *M. polymorpha*

Here, we reported that *MpSGF10B* is involved in the induction of reproductive organs in *M. polymorpha* (Figure 4C). In reproductive induction in liverworts, far-red light and circadian rhythms of long-day are key conditions, and these environmental stimuli are sensed and transduced by various signaling genes (Kohchi et al., 2021; Yamaoka et al., 2021). *MpBNB* acts as a key regulator of reproductive development (Yamaoka et al., 2018). Homologous modules are involved in light response and gametogenesis in *A. thaliana*, suggesting the conserved roles of these signaling modules among land-plant species (Kohchi et al., 2021; Yamaoka et al., 2021). Although further analyses are needed, the *MpSGF10B* homologs found in various plants might function in a conserved manner in reproductive development.

We found promoter activity of *MpSGF10B* in the following organs in *M. polymorpha:* bottom of gemma cups, surrounding areas of the meristem, and antheridia (Figures 2B,C). These expression patterns largely overlap with the accumulation patterns of auxin (Ishizaki et al., 2012). Auxin is proposed to control the development of various organs in *M. polymorpha*, including gemmae formation and the branching frequency of thalli (Kato et al., 2020). Notably, auxin-responsive genes were significantly repressed in gametophores, suggesting that attenuation of the auxin signal correlates with the transition to the reproductive phase in *M. polymorpha* (Flores-Sandoval et al., 2018). Additionally, recent studies showed that RALF1-FER inhibited root growth by induction of auxin biosynthesis in *A. thaliana* (Li et al., 2022), suggesting crosstalk between signaling peptides and phytohormones (Oh et al., 2018).

Taken together, MpSGF10B may interact with a partner(s) involved in auxin-regulated reproductive development. Despite the similarity in promoter activity between *MpSGF10B* and the auxin-responsive gene *GH3*, the *Mpsgf10b* mutant did not show any phenotypic changes, which were expected when perturbing the auxin signal. Thus, its contribution to auxin signaling might be small. Further analysis, including identifying MpSGF10B partners, could provide insights into signaling crosstalk between peptides and phytohormones in *M. polymorpha*.

## 5 Materials and Methods

### 5.1 Plant materials and growth conditions

*M. polymorpha* male or female accessions, Takaragaike-1 (MpTak-1) or Takaragaike-2 (MpTak-2), respectively, were used as the wild type in this study. Plants were axenically grown on half-strength Gamborg’s B5 medium pH 5.7 containing 1% agar and 1% sucrose at 22°C under continuous white light. For reproductive induction, 2-week-old plants were incubated under long-day conditions (light: 16 h, dark: 8 h) supplemented with continuous far-red light at 18°C.

### 5.2 RNA extraction and RT-PCR

Thalli or antheridiophores frozen in liquid nitrogen were grinded with a mortar and a pestle into fine powder. Total RNA was extracted from the tissue powder using the RNeasy Plant Mini Kit (Qiagen). cDNA was synthesized using ReverTra Ace reverse transcriptase (Toyobo). Amplification was performed with GoTaq DNA polymerase (Promega).

### 5.3 Plasmid construction

The primers used in this study are listed in Supplemental Table S3. For the promoter GUS constructs, approximately 5,000 bp upstream sequences flanked the start codons of the MpSGFs were amplified from MpTak-1 genomic DNA using PrimeSTAR GXL polymerase (Takara Bio) and cloned into a linearized pENTR1A vector using In-fusion (Takara Bio). Inserts were transferred into pMpGWB104 vectors (Ishizaki et al., 2015) using a Gateway LR Clonase II enzyme mix (Thermo Fisher Scientific). For the construction of transient expression assay, total RNA was extracted from MpTak-1 thalli and cDNA was synthesized. The CDSs of the MpSGFs were amplified from the cDNA using PrimeSTAR GXL polymerase. A DNA fragment of modified Venus (YFP-4 × Gly), the fluorescent protein with four Gly-residues at its C-terminus, was amplified from a gateway vector. Three fragments of MpSGFs CDS, YFP-4 × Gly CDS, and linearized pENTE1A backbone were fused using In-fusion. The entry vector was transferred to a binary vector, pGWB602 (Nakamura et al., 2010), with a Gateway LR Clonase II enzyme mix. For constitutive expression of MpSGF10B-YFP in *Marchantia*, DNA fragments of MpSGF10B-YFP were amplified from the destination vector mentioned above (MpSGF10B-Venus in pGWB602) as a template. The fragments were fused with the linearized pENTE1A using In-fusion. The entry vector was transferred to a binary vector pMpGWB103 (Ishizaki et al., 2015), with a Gateway LR Clonase II enzyme mix. For genome editing, guide RNAs were designed at two positions for each target gene using the CRISPR-Cas9 system (Sugano et al., 2018). Double-stranded DNAs corresponding to the guide RNA sequences were generated and inserted into AarI-digested pMpGE013 by ligation reaction using Mighty Mix (Takara Bio).

### 5.4 Generation of transgenic lines

*M. polymorpha* transformants were obtained by *Agrobacterium-mediated* transformation from regenerating thalli using the AgarTrap method (Tsuboyama and Kodama, 2018). Two wild-type accessions, MpTak-1 (male) and MpTak-2 (female), were used as genetic backgrounds. Transgenic plants were selected with the abovementioned medium and were supplemented with 10 μg/mL hygromycin or 0.1 μM chlorsulfuron. The transformants were clonally purified through gemmae generations. To examine the target loci of genome editing lines, genomic DNAs were extracted from TG1 plants, and then the PCR products were amplified using GoTaq DNA polymerase, followed by direct sequencing (Sugano and Nishihama, 2018).

### 5.5 GUS staining

Transgenic plants were fixed with 90% acetone at 4°C and then stained in GUS staining solution (100 mM sodium phosphate buffer pH 7.4, 0.5 mM potassium ferrocyanide, 0.5 mM potassium ferricyanide, 10 mM EDTA, 0.1% Triton X-100, and 0.5 mg/ml 5-bromo-4-chloro-3-indolyl-b-D-glucuronic acid) overnight at 37°C in the dark for GUS staining. The GUS-stained samples were cleared with 70% ethanol and chloral hydrate solution.

### 5.6 Transient expression assay in *N. benthamiana*

The *Agrobacterium tumefaciens* GV3101 strain was transformed with plasmids harboring proteins of interest (POIs) and grown in LB media containing spectinomycin (50 μg/mL). Cultures grown for about 24 h at 28°C with rotation at 220 rpm were used to transform *N. benthamiana* leaves. POI-containing bacteria were centrifuged for 15 min at 2,800 × *g* and resuspended in water to 4-fold dilution for infiltration. The cells were infiltrated into the abaxial side of *N. benthamiana* leaves. The infiltrated plants were incubated under dark conditions for 2 d. For imaging, the leaves were infiltrated with water (control) or 0.8 M D-mannitol solution (plasmolysis) using a 1 mL syringe, and leaf disks were harvested immediately.

### 5.7 Confocal fluorescent microscopy

The fluorescent images were inspected under a confocal laser scanning microscope (LSM780; Zeiss) using a 514- or 561-nm laser line. Fresh samples were mounted in water (control) or 0.8 M D-mannitol solution (plasmolysis) and imaged immediately.

### 5.8 Co-expression and GO enrichment analyses

Publicly available fastq data from RNA-Seq libraries were obtained from the sequence read archive in NCBI. All libraries used for analyses were performed in triplicate (except for antheridia duplicates). Fastq sequence data were mapped onto the reference genome of *M. polymorpha* (version 6.1) using STAR (Dobin et al., 2013), and the number of fragments was counted and normalized by transcripts per million (TPM) with RSEM (Li and Dewey, 2011). Accession numbers include: 11-day thalli (DRR050343, DRR050344, and DRR050345), Archegoniophore (DRR050351, DRR050352, and DRR050353), Antheridiophore, (DRR050346, DRR050347, and DRR050348), Antheridia (DRR050349, and DRR050350), apical cell (SRR1553294, SRR1553295, and SRR1553296), 13d-Sporophyte (SRR1553297, SRR1553298, and SRR1553299), Sporelings 0 h (SRR4450262, SRR4450261, and SRR4450260), 24hr-Sporeling (SRR4450266, SRR4450265, and SRR4450259), 48hr-Sporeling (SRR4450268, SRR4450264, and SRR4450263), 72hr-Sporeling (SRR4450267, SRR4450258, and SRR4450257), 96hr-Sporeling (SRR4450256, SRR4450255, and SRR4450254), thalli-mock (SRR5905100, SRR5905099, and SRR5905098), and 2,4-D 1h (SRR5905097, SRR5905092, and SRR5905091), plants infected with *P. palmivora* of 2 dpi (SRR7977547, SRR7977549, and SRR7977550), and mock-inoculated plants of 2 dpi (SRR8068335, SRR8068336, and SRR8068340) (Carella et al., 2018; Flores-Sandoval et al., 2018). Pearson’s correlation coefficients (PCCs) were calculated for the TPM of MpSGF10B and all genes using the R application. The top 1% (182 genes) with high PCC values were selected as co-expressed genes with MpSGF10B. These gene IDs were converted to those in assembly version 3 using the “convert ID” tool in MarpolBase (https://marchantia.info/). Then, GO enrichment analysis was performed using the Plant Transcriptional Regulatory Map (PlantRegMap, http://plantregmap.gao-lab.org/).

### 5.9 Statistical analysis

Statistical analyses were performed in R version 4.0.3. One-way ANOVA and one-tailed Dunnett’s *t*-test in package “multicomp” were used for multiple comparisons of means grouped by technical replicates.

## 7 Author Contributions

H. K. and T. S. conceived and designed the research in general; H. K. performed most of the experiments and analyzed the data; S. S. S. supported data analysis; K. T., Y. O., and T. M. supervised the experiments; H. K. and T. S. wrote the manuscript; All authors read, edited, and approved the manuscript.

## 9 Acknowledgments

We thank our colleagues for technical assistance, especially Kenta C. Moriya for culturing *M. polymorpha* and inducing reproductive organs, and Kazuaki Murakami for visualizing fluorescent proteins. We also thank Tsuyoshi Nakagawa (Shimane University, Japan) and Shoji Mano (National Institute for Basic Biology, Japan) for sharing the vectors. The authors would like to thank to Enago (www.enago.jp) for the English language review.

## 10 Supplementary Materials

**Supplementary Figure S1.** Multiple Sequence Alignments between MpSGFs and AtsORF.

**Supplementary Figure S2.** Relative expression levels of MpSGFs in various organs of *M. polymorpha*.

**Supplementary Figure S3**. Co-localization of MpSGF6 and MpSGF17 with chloroplasts.

**Supplementary Table S1.** A tabular of MpSGFs containing results of BLAST, annotations registered in the database (MarpolBase), existence of signal peptide sequence, and prediction of subcellular localization.

**Supplementary Table S2.** A list from GO enrichment analysis of co-expressed genes with each MpSGFs.

**Supplementary Table S3.** A list of primers used in this study.

